# Optical Genome Mapping of the human reference iPSC line KOLF2.1J reveals new smaller structural variants in neurodevelopmental genes

**DOI:** 10.1101/2024.10.17.618968

**Authors:** Madison James Yang, Kamilla Sedov, Max Y. Chen, Faria Zafar, Birgitt Schüle

## Abstract

The INDI consortium curated the KOLF2.1J human iPSC line to create a reference cell line for neurological disease modeling. However, despite careful assessments, two separate studies found using SNP arrays identified five structural variants (SVs) with sizes >100kbp. Two heterozygous SVs overlap the genes *JARID2, DTNBP1*, and *ASTN2*, raising concerns about KOLF2.1J’s suitability as a reference line. To investigate further, we screened KOLF2.1J for SVs smaller than 100kbp using optical genome mapping (OGM) to produce a high-resolution karyotype. OGM, validated by qPCR, indicated that one of the five known SVs contained a previously undetected overlap with *RYBP*. RYBP plays regulatory roles in neuronal differentiation, PAX6 expression and Notch signaling. Furthermore, OGM identified 11 SVs smaller than 100kbp, whose overlaps include the genes *PER2, CSMD1,* and *PALS1*. In summary, these mutations should be considered by researchers when using KOLF2.1J as a reference iPSC line for designing studies and experiments.

## Main Text

KOLF2.1J is a novel human induced pluripotent stem cell (hiPSC) line, derived from the parental HPSI0114i-kolf 2-C1 hiPSC line^1^. It was edited via CRISPR/Cas9 to repair *ARID2* and curated with the intent to serve as a reference cell line for modeling neurological diseases. Significant effort by the iPSC Neurodegenerative Disease Initiative (iNDI) consortium was devoted to quality control, ensuring that the KOLF2.1J line and its genetically engineered subclones were genetically stable, free of deleterious mutations, and capable of differentiating into neuronal cell types before public distribution^1^. Stability was assessed through G-banding karyotyping and genomic hybridization arrays. At the time of release, the line and its subclones were deemed free of structural variants (SVs) or other mutations that could affect their usability^1^.

However, two subsequent investigations, each employing high-density Illumina SNP arrays (Infinium GSA-24v3 BeadChip with 620,000 probes^2^ and Infinium Global Diversity Array with exonic coverage and 1.8 million probes^3^), identified five heterozygous structural variants (SVs) in KOLF2.1J, each more than 100 kbp in length. These included three heterozygous deletions and two heterozygous duplications, ranging in size from 132 to 766 kbp. Two SVs, affecting the genes *JARID2* and *DTNBP1* (6p22.3 deletion), and *ASTN2* (9q33.1 deletion), have been associated with neurodevelopmental disorders and showed 50% reduced gene expression in iPSCs and neural progenitors. In contrast, while the 3p14 duplication affected several genes, none were classified as deleterious. Additionally, the 18q22 duplication and the 3p13 deletion did not overlap any known genes.

To search for undetected pathogenic mutations in KOLF2.1J, we focused on identifying structural variants (SVs) smaller than 100 kb, utilizing optical genome mapping (OGM) in the Bionano Genomics platform. This technology labels ultra-high molecular weight DNA with a 6-nucleotide marker and images the labeled DNA patterns to produce a high-resolution karyotype. OGM can detect all classes of structural variants, with a resolution down to 500 base pairs (bp) and a 5% variant allele frequency. OGM has a label density (14-17 labels/kbp), providing enhanced positional accuracy over SNP arrays.

Our objective was to evaluate the reliability of OGM by testing it against the known SVs in KOLF2.1J and determining its ability to detect smaller SVs in this iPSC line. Using OGM, we confirmed the presence of all five previously identified SVs^2^ (Supplemental Figure 1, Supplemental Table 1). We also made two critical discoveries using OGM: 1) An overlapping gene in the 3p13.2 (143.8 kbp) deletion; and 2) Eleven additional smaller heterozygous SVs in KOLF2.1J that were not previously described, ranging from 4.8 kbp to 80.9 kbp in size (Supplemental Figure 2, Supplemental Table 1).

Previously, the Illumina SNP arrays did not find any overlap with other genes in the 3p13.2 (143.8 kbp) deletion. However, our OGM data pinpoint that the 3p13.2 deletion overlaps the *RING1* and *YY1* binding protein (*RYBP*) gene, deleting exons 3-4. OGM analysis shows that the 3p13.2 deletion is larger than previously described in the two prior studies^2, 3^, with the 5’-end of 72,425,487 lying within the *RYBP* gene (T2Tchm13v2.0, Chr3:72,413,401 - 72,488,070) (Supplemental Figure 3). We further verified by qPCR a ∼50% reduction in relative gene expression for *RYBP* in KOLF2.1J iPSCs (Supplemental Figure 4), implicating *RYBP*’s inclusion in the deleted region.

RYBP is a key component of the Polycomb repressive complex 1 (PRC1) and mediates histone H2A ubiquitination (H2AK119ub1), playing a crucial role in neuronal development^4^. Its haploinsuffiency leads to impaired neuronal differentiation, with an expansion of neural progenitors and a failure to mature into functional neurons^5^. This is linked to altered retinoic acid (RA) signaling and differential PAX6 expression^6^; a delicate PAX6 gradient is critical to proper hindbrain patterning *in vivo*^7^. We found that PAX6 shows significantly decreased expression in KOLF2.1J throughout neuronal differentiation relative to differentiated control iPSCs, prior to differentiation and at days 21 and 31 in differentiation (Supplemental Figure 5). RYBP depletion also reduces dendritic complexity and inhibits Notch signaling, crucial for neurogenesis^8^. These findings suggest that RYBP functions as a transcriptional repressor that regulates pathways essential for neurodevelopment, including axon guidance and synaptic signaling.

Furthermore, with OGM, we detected 11 additional smaller heterozygous SVs in KOLF2.1J, ranging from 4.8 kbp to 80.9 kbp in size, that were not previously described due to the limitations of standard SNP arrays (Supplemental Table 1). Six of these SVs overlap exons of genes (Supplemental Table 1), and three SVs could be potentially deleterious and are implicated in neurological disorders or neuronal development. We curated the genes *in silico* based on their gnomAD^9^ pLi and gnomAD LOEUF scores, ClinVar^10^ assessment, and their effects in mice when deleted on both alleles^11^ (Supplemental Table 3).

A 2q37.3 insertion overlaps *PER2*, a well-studied, ubiquitously expressed gene critical to circadian rhythm, and greatly affected by gene dosage changes^12^. We found a reduced relative expression level of 0.59 for *PER2*. An 8p23.2 deletion overlaps *CSMD1*, a complement system component where loss-of-function causes impaired coordination of neuronal differentiation^13^. *CSMD1* polymorphisms are also a significant risk factor for Parkinson’s disease^14^. Finally, a 14q23 deletion overlaps *PALS1*, a gene involved in cell polarity and Schwann cell recruitment^15^.

OGM represents a paradigm shift in detecting SVs for iPSC quality control. Our findings indicate that the KOLF2.1J line contains SVs that researchers using this line should be aware of. A key advantage of OGM is its precise size resolution, which significantly reduces false negatives, such as those related to the 3p13.2 RYBP deletion. For rigorous quality control, detecting these SVs in iPSCs is essential, both immediately after nuclear reprogramming and throughout regular passaging, to identify potential in vitro artifacts and expansion of mutated clones. Traditional SNP arrays lack the range and resolution necessary for detecting smaller SVs, in contrast to OGM. Ultimately, OGM addresses a critical technological gap, identifying SVs at low variant allele frequencies and enabling the replacement of iPSC lines with mutations in genes crucial to context-specific research.

## Acknowledgements

The study was conducted with departmental start-up funds and philanthropic support (B.S.).

## Declaration of Interests

The authors declare no competing interests.

## Supplemental Material

### Supplemental Figures

**Supplemental Figure 1:**
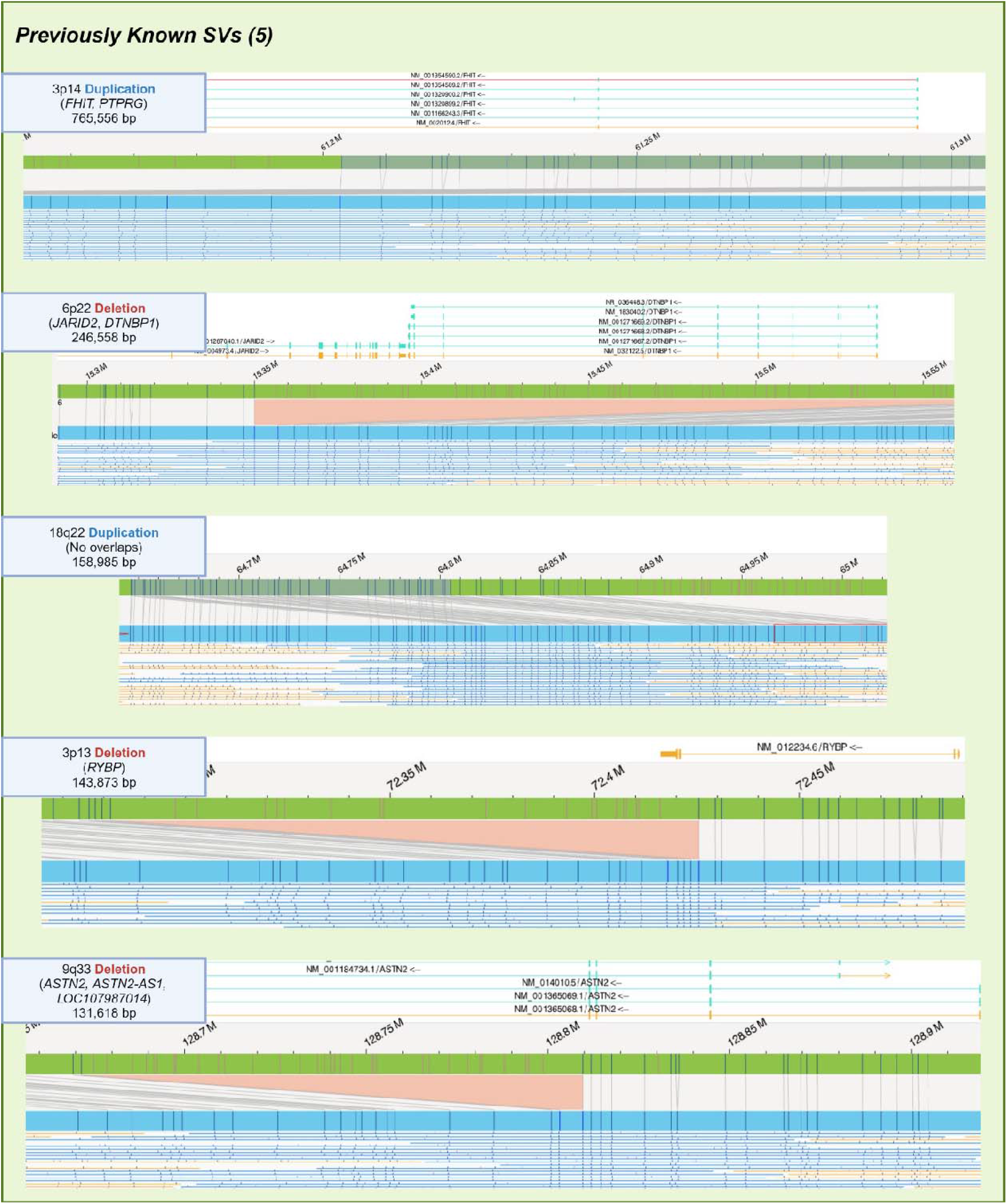
OGM overview of the five previously known SVs. SNP array was able to identify these five deletions and duplications, which were also identified with OGM analysis. OGM displays the SVs as a change between the reference (green) and variant assembly (blue), with individual supporting reads shown below. Figure created with BioRender.com.

**Supplemental Figure 2:**
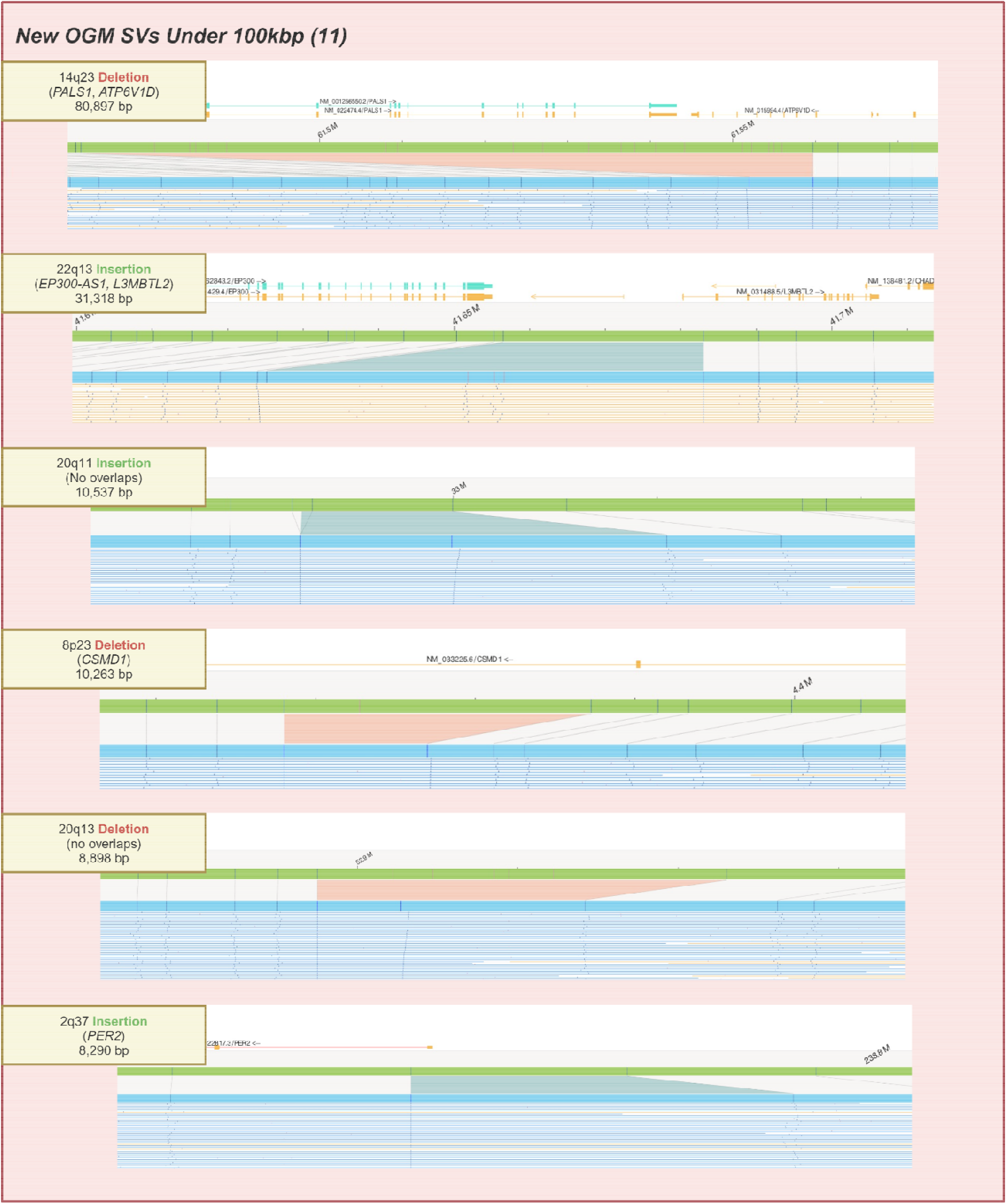
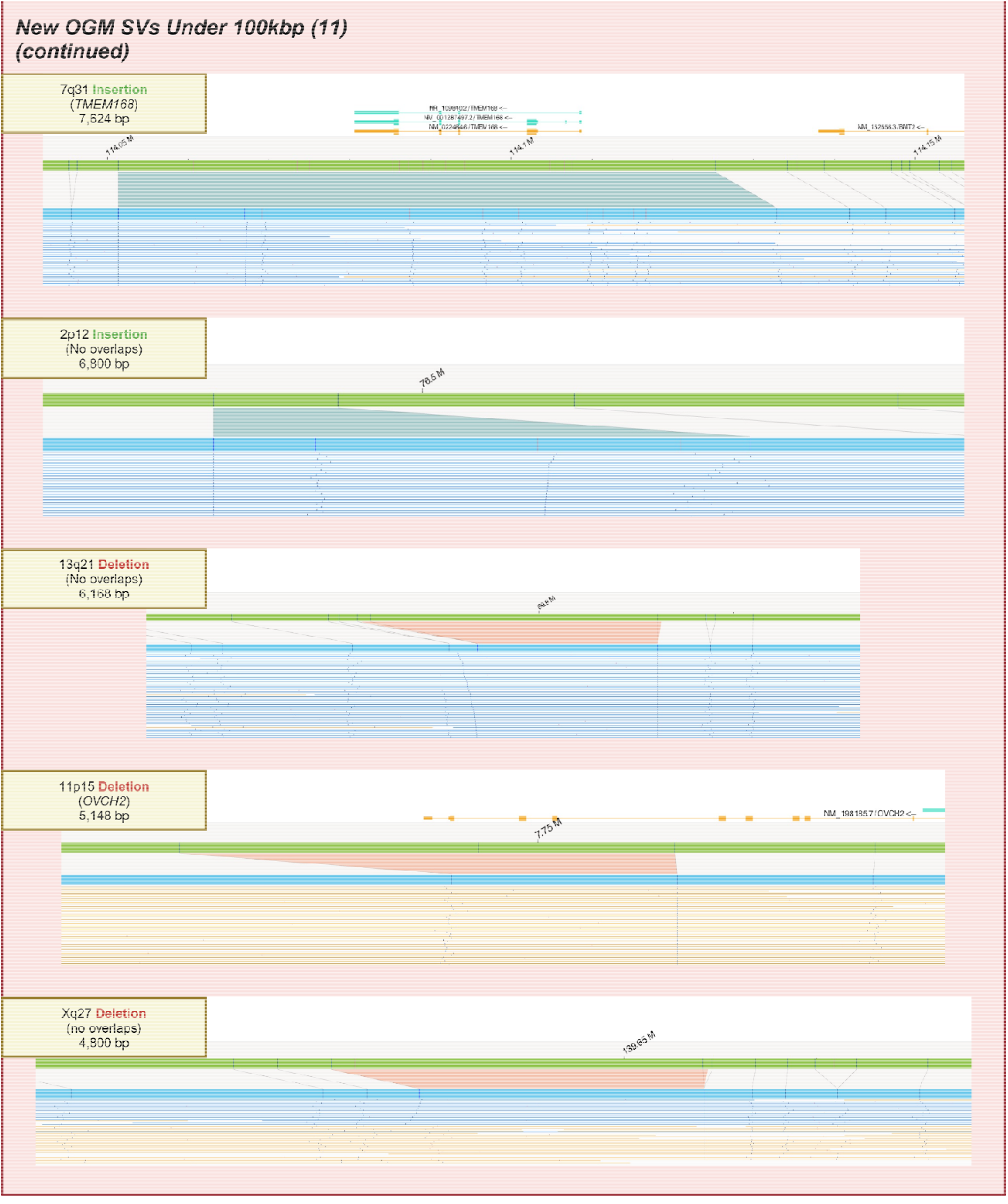
OGM overview of the 11 OGM-exclusive SVs. These 11 SVs, comprising a mix of insertions and deletions, all have lengths under 100kbp and were identified with OGM analysis alone. Figure created with BioRender.com.

**Supplemental Figure 3:**
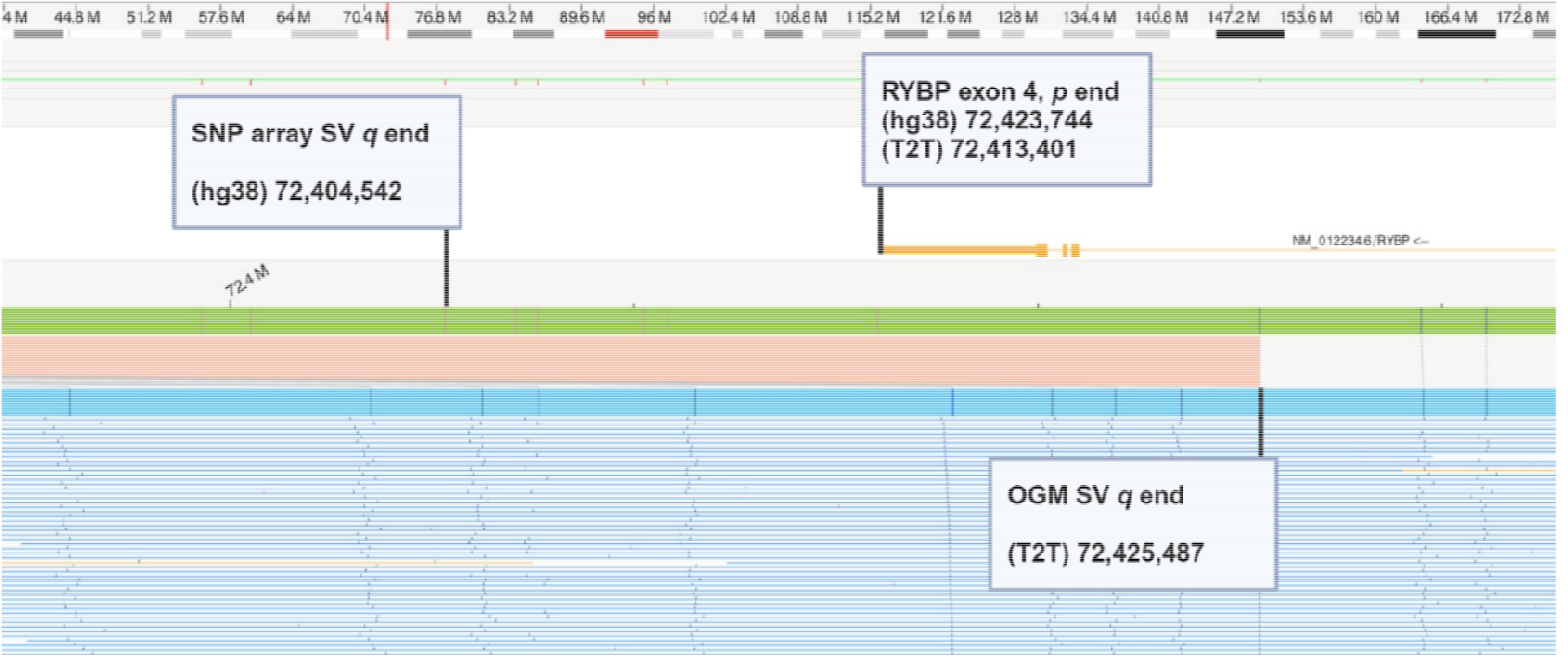
In-detail overview of 3p13 deletion. Inspection of chromosome 3 deletion overlapping the *RYBP* gene. Previously, SNP array identified a smaller deletion on chromosome 3 with no overlap, but the enhanced SV resolution from OGM shows that the SV overlaps *RYBP*.

**Supplemental Figure 4:**
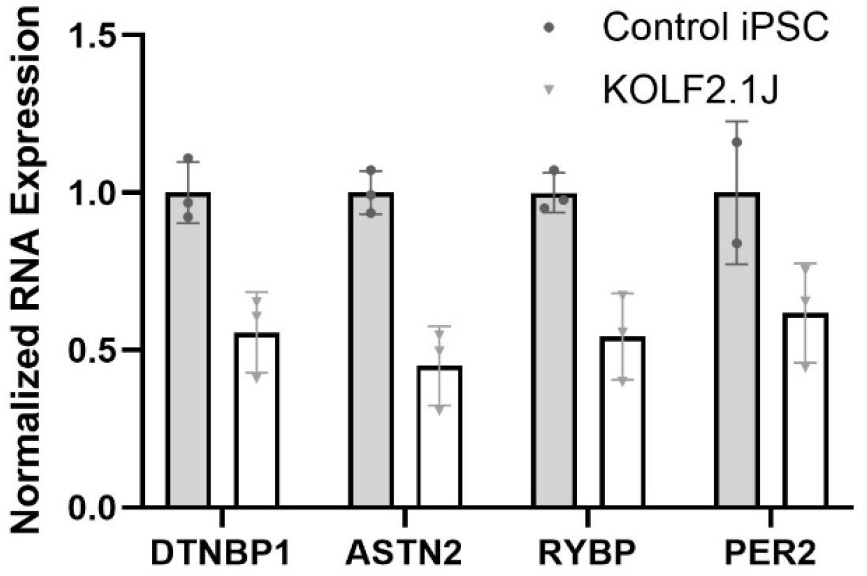
qPCR relative fold change for genes affected by SVs. Displayed are the mRNA expression levels for four SV-affected KOLF genes, two previously identified and two newly identified with OGM. Values are normalized relative to the housekeeping gene GAPDH, then normalized to values for our in-house control line.

**Supplemental Figure 5:**
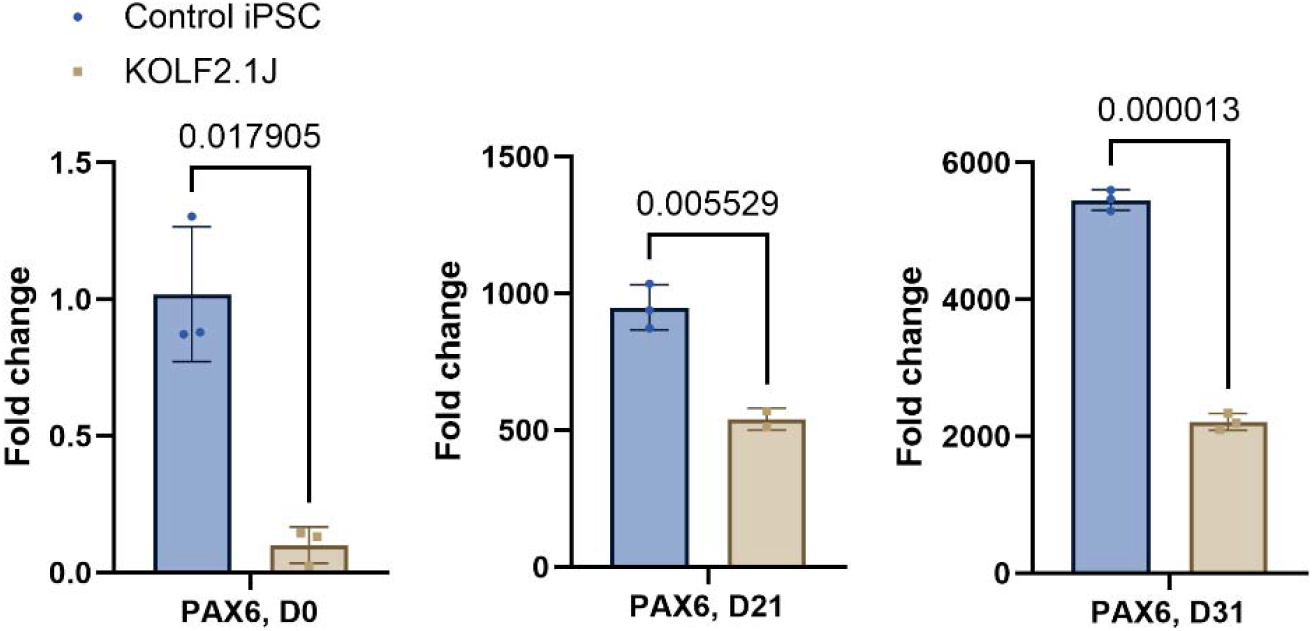
qPCR relative fold change for PAX6 in KOLF2.1J at different stages in differentiation. At the D0 (day 0, pre-differentiation), D21, and D31 time points in their respective neuronal differentiation, the fold change (normalized to controls at the iPSC/D0 stage) for PAX6 is significantly lower in the KOLF2.1J line relative to controls.

### Supplemental Tables

**Supplemental Table 1:** SV data collected for KOLF2.1J. All SVs found that have a 0% presence in the Bionano control, sorted by size. The five SVs previously known are highlighted in gray. Genes identified as potentially deleterious are labeled red and italicized.

| Chromosome/<br>Cytoband | Start | End | SV Type | Length<br>(BP) | Allele<br>Fraction | Molecule<br>Support | Affected Genes/ Transcripts |
| --- | --- | --- | --- | --- | --- | --- | --- |
| 3p14.2 | 61202985.5 | 61968541 | Duplication | 765556 | 1 | 107 | FHIT, PTPRG |
| 6p22.3 | 15350000 | 15603593 | Deletion | 246558 | 0.49 | 93 | *JARID2, DTNBP1* |
| 18q22.1 | 64646213 | 64805198 | Duplication | 158985 | 0.47 | 121 | - |
| 3p13 | 72281614 | 72425487 | Deletion | 143873 | 0.52 | 121 | *RYBP* |
| 9q33.1 | 128671603 | 128809609 | Deletion | 131618 | 0.51 | 100 | *ASTN2*, ASTN2-AS1,<br>LOC107987014 |
| 14q23.3 | 61471120 | 61559693 | Deletion | 80897 | 0.45 | 98 | *PALS1*, ATP6V1D |
| 22q13.2 | 41656483 | 41683001 | Insertion | 31318 | 0.53 | 88 | EP300-AS1, L3MBTL2 |
| 20q11.21 | 32992556.6 | 32999995.4 | Insertion | 10537 | 0.52 | 128 | - |
| 8p23.2 | 4368040 | 4387258.5 | Deletion | 10263 | 0.57 | 108 | *CSMD1* |
| 20q13.13 | 52897510 | 52935008 | Deletion | 8898 | 0.51 | 19 | - |
| 2q37.3 | 238777272 | 238788080 | Insertion | 8290 | 0.5 | 103 | *PER2* |
| 7q31.1 | 114051331 | 114125280 | Insertion | 7624 | 0.42 | 103 | TMEM168 |
| 2p12 | 76496551 | 76498608 | Insertion | 6800 | 0.53 | 114 | - |
| 13q21.33 | 69790841.5 | 69806289.6 | Deletion | 6168 | 0.48 | 99 | - |
| 11p15.4 | 7743180 | 7752613 | Deletion | 5148 | 0.43 | 43 | OVCH2 |
| Xq27.2 | 139634768.9 | 139654349.6 | Deletion | 4800 | 0.76 | 25 | - |

**Supplemental Table 2:** Quality control metrics at each stage of OGM processing. These metrics cover the quantitative measurements taken in the extraction, labeling, and read generation stages of OGM. Metrics for KOLF2.1J fall within the ideal cutoffs for each of the required QC limits.

| QC Metric<br>Category | Post-Elute<br>gDNA<br>Conc. | Post-Label<br>gDNA<br>Conc. | CV<br>($\sigma / \bar{X}$)<br>of Conc. | N50><br>150kbp | N50><br>20kbp | Coverage | Map<br>Rate | Label Density |
| --- | --- | --- | --- | --- | --- | --- | --- | --- |
| **Ideal**<br>**Range** | 40-150 ng/μL | 4-16 ng/μL | <0.3 | >230kbp | >150kbp | >350X | >80% | 14-17 labels/kbp |
| **KOLF2.1J**<br>**Data** | 122.3 ng/μL | 6.985 ng/μL | 0.04 | 241.1kbp | 178.13kbp | 425.85X | 91.6% | 16.38 labels/kbp |

**Supplemental Table 3:**
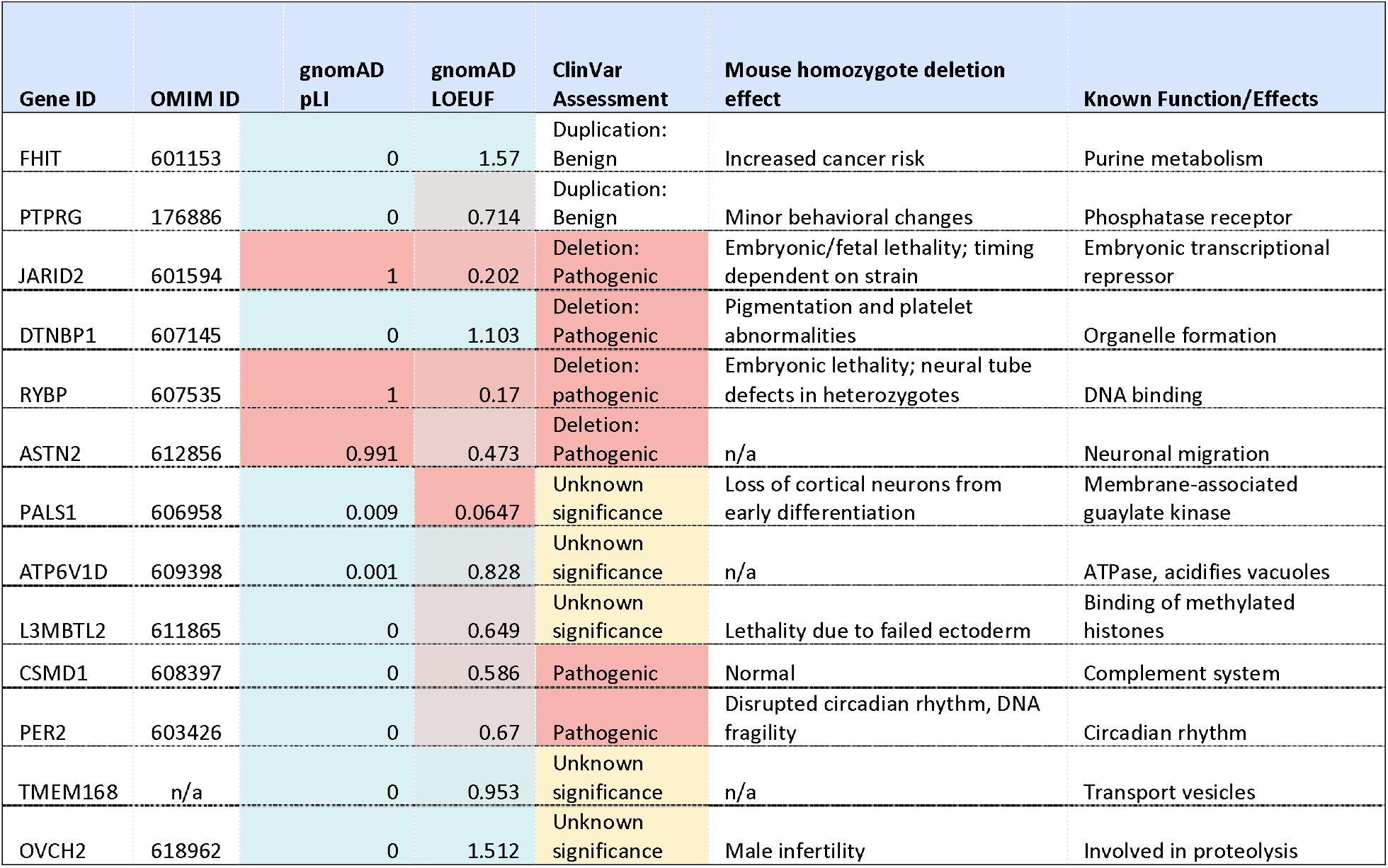
Table of genes found in KOLF2.1J SVs and their potential deleterious effect. Known information of each gene involved in SVs found in KOLF2.1J, the SV effect, and their pathological significance. The most important criterion is the gnomAD LOEUF; a value below 0.63 is considered potentially pathogenic (although pathogenic SVs exist that are above this cutoff). Note that the transcripts ASTN2-AS1, LOC107987014, and EP300-AS1 are excluded as they are noncoding sequences for which there is no relevant data.

**Supplemental Table 4:**
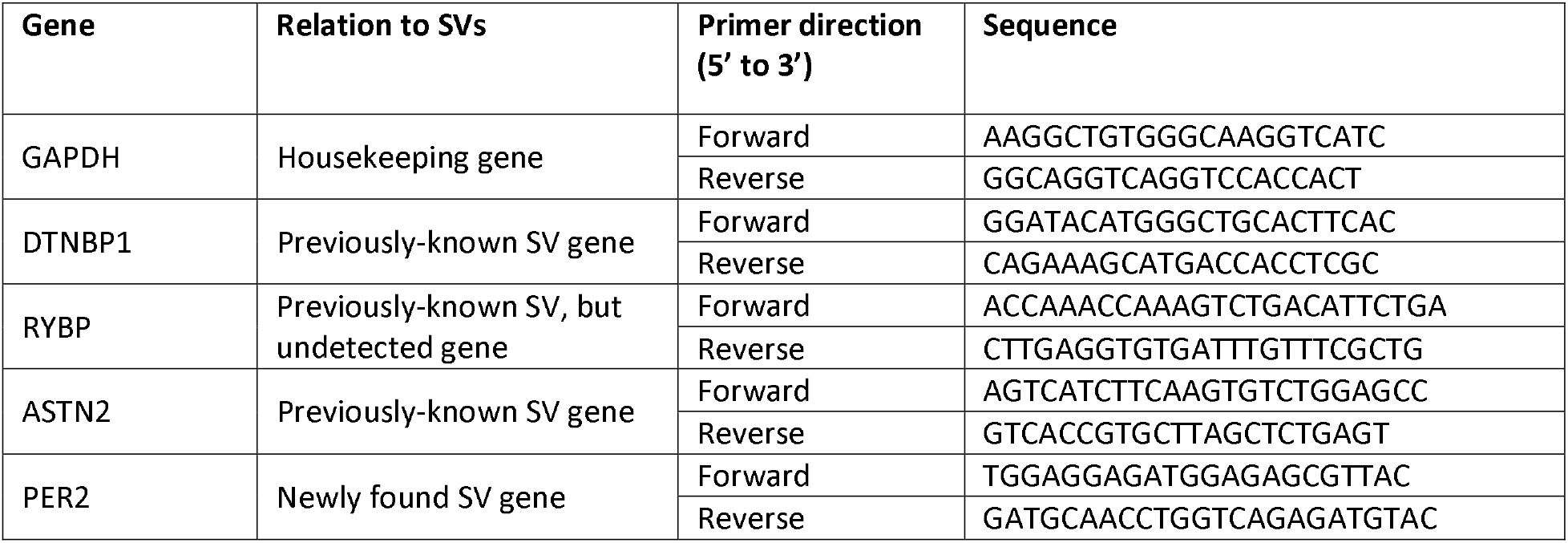
List of primers used for KOLF2.1J SYBR Green qPCR assay. These five primer pairs were designed to flank five genes of interest: one housekeeping control to measure relative expression against, two known heterozygous deletions (previously confirmed by SNP array), and two of the newly-discovered SVs in KOLF2.1J, also heterozygous deletions.

**Supplemental Table 5:** Sybr Green qPCR reaction cycle scheme.

| Steps | Temperature | Duration | Cycles |
| --- | --- | --- | --- |
| UDG activation | 50 °C | 2 min | 1 |
| Dual-Lock™ Taq DNA polymerase | 95 °C | 2 min | 1 |
| Denaturation | 95 °C | 15 s | 40 |
| Annealing/ extend | 60 °C | 1 min |  |

**Supplemental Table 6:** Sybr Green dissociation cycle scheme.

| Step | Ramp rate | Temperature | Time |
| --- | --- | --- | --- |
| Denaturation | 1.6 °C/s | 95 °C | 15 s |
| Annealing | 1.6 °C/ s | 60 °C | 1 min |
| Dissociation | 0.15 °C/ s | 95 °C | 15 s |

### Supplemental Methods

#### Bionano optical genome mapping

Ultra-high molecular weight DNA was extracted from the cell pellets (1-1.5×10^6 cell/pellet) using the *Bionano Prep SP-G2 Blood & Cell Culture DNA Isolation Kit (Bionano Genomics, PN 80060)* as according to Bionano-issued protocol CG-00004. Cell pellets were carefully resuspended, lysed, and depleted of proteins with thermolabile Proteinase K, which was then deactivated with a heated-lid thermocycler. DNA was bound to NanoBind Discs (PacBio, included in kit) and cleaned with successive washes, then subsequently eluted and incubated overnight to ensure sample concentration homogeneity. Eluted DNA samples were then labeled with the enzyme DLE-1, which applies the DL-Green fluorescent label at the palindromic sequence CTTAAG, and counterstained blue. Labeled gDNA samples were loaded into flowcell chips (Bionano Genomics, PN 20440) and imaged through the Bionano Saphyr instrument, with a flowthrough target of 1500 GBps (corresponding to approximately 400x target coverage). This process images the DNA into molecule reads by passing them through microfluidic channels and directly observing label patterns from laser photography. Between the extraction, labeling, read generation, and analysis steps, a series of quality control measurements were taken, to ensure fidelity of reads gathered (Supplemental Table 2). DNA concentrations were measured with a Qubit fluorometer.

SV alignments were generated from collected reads using the Rare Variant Assembly workflow of Bionano Access version 1.8.1.1, using T2T-CMH13v2.0 as the selected reference alignment as it is gapless^16^. SVs generated this way were filtered based on the following criteria: valid SVs contain at least five molecule reads that completely span all markers in the SV assembly, and are not present in the Bionano control database, a repository of SVs from 285 healthy individuals. Genes found to overlap SVs identified this way were rated for potential pathogenicity based on the on the following databases (Supplemental Table 3): 1. OMIM (Online Mendelian Inheritance in Man^17^) was used for determining the existence of known pathogenic phenotypes. 2. Genes were scored through the genome-agnostic database gnomAD^9^, which assigns a normalized LOEUF (loss-of-function observed/expected upper bound fraction) score to each gene (values under 1 indicate genomic constraint, i.e. purifying selection). A low LOEUF directly implies strong susceptibility to mutational strain. Therefore, we chose a maximum cutoff of 0.63 for potentially deleterious genes, being the mean LOEUF of essential genes. 3. ClinVar^10^ was used to search for the existence of known variants and their pathogenic status. Samples were assessed based on the presence of pathogenic variant that share copy number change type. Direct confirmation of effect through these databases is not possible for all genes as some genes lack information of prior study and/or lack of reference ClinVar variation. 4. Additional data was used from the JAX Mouse Genome Database^11^, listing the homozygous effect of deletions of equivalent homologs in mice, to supplement judgements.

#### SYBR Green expression analysis

qPCR was performed to verify the existence of changes in genes identified this way in KOLF2.1J. Using a previously established 96 -well SYBR Green qPCR expression array^18^, iPSCs from KOLF2.1J were characterized. iPSCs were dissociated with ReLesR and ∼2,500,000 cells were pelleted for 5 min by centrifugation (5,000 g at 4°C). The pellet was washed with 1 ml of PBS and again centrifuged at 5,000 g at 4°C. RNA was then extracted using the Purelink™□ RNA Mini Kit. The RNA concentration was quantified by NanoDrop and 1 μg of RNA was treated with DNase I for 15 min at RT to eliminate contamination with genomic DNA. After inactivating DNase I for 3 min at 65°C for 3 min, HighCapacity cDNA Reverse Transcription kit was used for cDNA reverse transcription. Negative control was generated by adding nuclease-free water instead of RNA. For the reverse transcriptase control, RNA was added but not the reverse transcriptase. Each cDNA sample was diluted 1:5 with nuclease-free H_2_O for a final concentration of 10 ng/μL and stored at -20°C. The day before the SYBR Green assay, both forward and reverse primers for the genes of interest (Supplemental Table 4) were pre-plated in triplicates at a final concentration of 300 nM each into a 384-well of an optical PCR plate. Additional wells with GADPH (housekeeping gene) primers were included for the negative and reverse transcriptase control. After plating the primers, the 384 well-plate was centrifuged at 2,000 rpm for 2 min. To dry out the primers, the plate was kept in a box at RT for 12 h. Then the reaction mix (2.5 μL of PowerUp SYBR Green Master Mix, 1.5 μL of nuclease-free water, and 1 μL of 10ng/μL sample cDNA) was pipetted to the wells that were pre-coated with primers, and the plate was sealed with optical adhesive film. Following another centrifugation step at 2,000 rpm for 2 min, the SYBR Green reaction was run using a QuantStudio 6 Flex. The cycling conditions indicated in Supplemental Table 5 were used to amplify cDNA, while the conditions in Supplemental Table 6 were used to assess the melting curve of the PCR product. The cycle threshold (Ct) values were used to calculate the fold change (2^-ΔΔCt^) as a measure of relative gene expression. First, the mean Ct value was calculated for each gene. The mean Ct values were used to determine ΔCt by ΔCt = mean Ct target gene - mean Ct housekeeping gene (GAPDH). Finally, fold change was by normalizing the Ct values to our in-house control iPSC line 4using ΔΔCt = ΔCt KOLF2.1J – ΔCt control, and the fold change was obtained by 2^-ΔΔCt^.

## Notes

### Competing Interest Statement

The authors have declared no competing interest.

